# Postnatal Expansion, Maturation and Functionality of MR1T Cells in Humans

**DOI:** 10.1101/2019.12.20.882746

**Authors:** Gwendolyn M. Swarbrick, Anele Gela, Meghan E. Cansler, Megan D. Null, Rowan B. Duncan, Elisa Nemes, Muki Shey, Mary Nsereko, Harriet Mayanja-Kizza, Sarah Kiguli, Jeffrey Koh, Willem A. Hanekom, Mark Hatherill, Christina Lancioni, David M. Lewinsohn, Thomas J. Scriba, Deborah A. Lewinsohn

## Abstract

MR1-restricted T (MR1T) cells are defined by their recognition of metabolite antigens presented by the monomorphic MHC class 1-related molecule, MR1, the most highly conserved MHC class I related molecule in mammalian species. Mucosal-associated invariant T (MAIT) cells are the predominant subset of MR1T cells expressing an invariant TCR α-chain, TRAV1-2. These cells comprise a T cell subset that recognizes and mediates host immune responses to a broad array of microbial pathogens, including *Mycobacterium tuberculosis*. Here, we sought to characterize development of circulating human MR1T cells as defined by MR1-5-OP-RU tetramer labelling and of the TRAV1-2^+^ MAIT cells defined by expression of TRAV1-2 and high expression of CD26 and CD161 (TRAV1-2^+^CD161^++^CD26^++^ cells). We analysed postnatal expansion, maturation and functionality of peripheral blood MR1-5-OP-RU tetramer^+^ MR1T cells in cohorts from three different geographic settings with different tuberculosis (TB) vaccination practices, levels of exposure to and infection with *M. tuberculosis.* Early after birth, frequencies of MR1-5-OP-RU tetramer^+^ MR1T cells increased rapidly by several fold. This coincided with the transition from a predominantly CD4^+^ and TRAV1-2^−^ population in neonates, to a predominantly TRAV1-2^+^CD161^++^CD26^++^ CD8^+^ population. We also observed that tetramer^+^ MR1T cells that expressed TNF upon mycobacterial stimulation were very low in neonates, but increased ∼10-fold in the first year of life. These functional MR1T cells in all age groups were MR1-5-OP-RU tetramer^+^TRAV1-2^+^ and highly expressed CD161 and CD26, markers that appeared to signal phenotypic and functional maturation of this cell subset. This age-associated maturation was also marked by the loss of naïve T cell markers on tetramer^+^ TRAV1-2^+^ MR1T cells more rapidly than tetramer^+^TRAV1-2^−^ MR1T cells and non-MR1T cells. These data suggest that neonates have infrequent populations of MR1T cells with diverse phenotypic attributes; and that exposure to the environment rapidly and preferentially expands the MR1-5-OP-RU tetramer^+^TRAV1-2^+^ population of MR1T cells, which becomes the predominant population of functional MR1T cells early during childhood.

## Introduction

MR1-restricted T (MR1T) cells are T-cells that display a highly restricted T-cell-receptor (TCR) repertoire and exhibit innate-like functions similar to those described for NKT cells. MR1T cells are defined by their recognition of metabolite antigens presented by the monomorphic MHC class 1-related molecule, MR1, the most highly conserved MHC class I related molecule in mammalian species (Harriff et al., 2018; Godfrey et al., 2019). The TRAV1-2^+^ MR1T cells are known as Mucosal Associate Invariant T cells (MAIT) and this subset express an invariant TCR α-chain (TRAV1.2-TRAJ33/20/12 in humans; TRAV1-TRAJ33 in mice) with a CDR3 of constant length paired with a limited number of Vβ segments (Vβ2/13 in humans; Vβ6/8 in mice) (Treiner et al., 2003; Martin et al., 2009; Reantragoon et al., 2013; Lepore et al., 2014). MR1T cells have been defined by different methods. Prior to availability of the MR1 tetramer, MR1T cells were phenotypically defined as the predominant MAIT cell subset, by expression of TRAV1-2 and high level expression of CD161 (Walker et al., 2012; Fergusson et al., 2014; Sharma et al., 2015) and CD26 (Sharma et al., 2015). Now MR1 tetramers can also be more broadly used to define MR1T cells. More specifically, the MR1-5-OP-RU tetramer, while not universally staining all MR1T cells, stains the vast majority of MR1 tetramer staining T cells and has been widely used to define this cell population (reviewed in Godfrey et al., 2019). In peripheral blood of adults, populations of MR1-5-OP-RU tetramer^+^ MR1T cells are highly concordant with populations of TRAV1-2^+^CD161^++^CD26^++^ phenotypically defined MAIT cells (Meermeier et al., 2016). As we will show in this paper, the MR1-5-OP-RU tetramer allows detection of a much broader repertoire of MR1T cells that are not only restricted to TRAV1-2^+^ MAIT cells. Moreover, we show that in neonates, MR1-5-OP-RU tetramer^+^ MR1T cells and TRAV1-2^+^CD161^++^CD26^++^ T cells are much less concordant populations.

MR1T cell development is a stepwise process, with an intra-thymic selection followed by peripheral expansion. Using the MR1-5-OP-RU tetramer, Koay *et al* reported a three-stage development pathway for mouse and human MR1T cell populations, where three populations of human MR1T precursors were defined based on differential expression of CD27 and CD161. The initial two stages of MR1-5-OP-RU tetramer^+^TRAV1-2^+^CD161^−^ cells, which were either CD27^−^ or CD27^+^, occurred in the thymus and these cells were low in the periphery. The more mature MR1-5-OP-RU tetramer^+^TRAV1-2^+^CD161^+^CD27^+^ cells, designated as stage 3, have been shown to represent only a small fraction of T cells in the thymus and neonates but form a more abundant population in adult blood (Martin et al., 2009; Dusseaux et al., 2011; Koay et al., 2016). Studies utilizing TRAV1-2 and high level CD161 expression (CD8^+^TRAV1-2^+^CD161^++^ cells) to define MAIT cells, similarly demonstrated low numbers of these cells in the neonates, which expand postnatally (Walker et al., 2012; Fergusson et al., 2014). Such studies indicate that MR1T cell thymopoeisis is complemented by an important postnatal peripheral expansion and maturation, which was recently shown to be dependent upon acquisition of the intestinal microbiome (Constantinides et al., 2019; Legoux et al., 2019).

Studies that define MAIT cells by phenotypic characteristics or by MR1 5-OP-RU staining demonstrate that these cells are activated via both TCR-dependent and TCR-independent mechanisms. Regarding TCR-dependent activation, MAIT cells recognize microbial derived riboflavin (Vitamin B2) biosynthesis intermediate derivatives, such as 5-OP-RU (5-(2-oxopropylideneamino)-6-d-ribitylaminouracil) and a broad spectrum of microbes that synthesize riboflavin, including *S. aureus, E. coli, and C. albicans* (Gold et al., 2010; Kjer-Nielsen et al., 2012; Gold et al., 2013; Corbett et al., 2014; Meermeier et al., 2016). While 5-OP-RU was the first activating MR1 ligand defined, several other ligands have subsequently been defined, including additional riboflavin-derived ligands, and both stimulatory and inhibitory ligands (Eckle et al., 2014; Gherardin et al., 2016; Meermeier et al., 2016; Harriff et al., 2018). In addition, cytokines, primarily IL-12, IL-18, and IL-15, act synergistically to activate MAIT cells, both by lowering the threshold of TCR activation (Ussher et al., 2014; Sattler et al., 2015; Slichter et al., 2016) and/or inducing MR1-independent activation (Fergusson et al., 2014; Ussher et al., 2014; Sattler et al., 2015; Slichter et al., 2016; van Wilgenburg et al., 2016; Suliman et al., 2019). These TCR-independent activation mechanisms are hypothesized to explain how MAIT cells may participate in host defense to viruses and bacteria that do not synthesize riboflavin (Ussher et al., 2014; van Wilgenburg et al., 2016). Once activated, MAIT cells display immediate effector function by secreting inflammatory cytokines and cytotoxic molecules, which can kill infected cells (Gold et al., 2010; Dusseaux et al., 2011; Le Bourhis et al., 2013; Kurioka et al., 2015). In a mouse model of pulmonary legionella infection, MAIT cells afforded enhanced protection in an MR1-dependent manner, mediated by the effector molecules IFN-γ and GM-CSF (Wang et al., 2018). MAIT cells have also been shown to facilitate control of *Klebsiella pneumoniae, Mycobacterium bovis* BCG, and *Francicella tularensis* infections (Georgel et al., 2011; Meierovics et al., 2013; Sakala et al., 2015; Meierovics and Cowley, 2016). Many of the bacterial pathogens that are important causes of infant morbidity and mortality, including *S. aureus, S. pneumoniae*, and enteric pathogens (e.g. *E. coli, Klebsiella, Salmonella*), are recognized by MAIT cells. Moreover, MAIT cells may contribute to viral immunity and hence host defense to viral pneumonia, which represents the major cause of morbidity and mortality worldwide (Zar and Ferkol, 2014). MAIT cells possess immediate innate-like, effector functions, and can therefore mediate immune functions before priming, clonal expansion and maturation of antigen-specific MHC-restricted T cells (Gold et al., 2013; Hartmann et al., 2018; Godfrey et al., 2019). Therefore, MAIT cells could play an important role in host defense to a variety of pathogens, especially in infancy.

MAIT cells are known to also recognize and respond to *Mycobacterium tuberculosis*, the leading global cause of mortality from a single infectious agent. Infants and young children suffer disproportionately from disseminated and severe forms of tuberculosis (TB) disease and are significantly more susceptible to disease than adults (Marais et al., 2004a; Marais et al., 2004b; Marais et al., 2006). Here, we sought to characterize development of MR1T cells (defined by MR1-5-OP-RU-tetramer staining) by analyzing postnatal expansion, maturation and changes in phenotype and functionality in humans. We compared MR1T cell development in cohorts from three different settings with differing TB vaccination practices and levels of exposure and infection with *M. tuberculosis.*

## Results

### Human participants

We enrolled nine cohorts across three different countries to study MR1T cell development (Table 1). The cohorts in South Africa (SA) were neonates (cord blood, *n* = 10), infants (aged 10 weeks, BCG vaccinated at birth, *n* = 10) and adolescents [aged 14-18 years, BCG vaccinated at birth, QuantiFERON-TB Gold (QFT)-negative, *n* = 10]. The cohorts enrolled in the United States (US) were neonates (cord blood, *n* = 10), infants (aged 9-15 months, BCG unvaccinated, *n* = 10) and adults (aged 29-60 years, *n* = 10). The cohorts in Uganda (UG) were infants [2-24 months old, BCG vaccinated at birth +/−, tuberculin skin test (TST) +/−, *n* = 36], children (24 - 60 months, BCG vaccinated at birth +/−, TST +/−, *n* = 23) and adults (aged 18-60 years, BCG vaccinated at birth +/−, TST +/−, *n* = 91).

**Table 1.** Study Participants.

| *Country* | *Cohort* | *number* | *Age* | *BCG vaccination* | *QFT results* | *TST results* |
| --- | --- | --- | --- | --- | --- | --- |
| ***South Africa*** | Neonates | 10 | 0 (cord blood) | - | ND | ND |
|  | Infants | 10 | 10 weeks | 100% | ND | ND |
|  | Adolescents | 10 | 14-18 years | 100% | 100% negative | ND |
| ***USA*** | Neonates | 10 | 0 (cord blood) | - | ND | ND |
|  | Infants | 10 | 9-15 months | - | ND | ND |
|  | Adults | 10 | 29-60 years | - | ND* | ND* |
| ***Uganda*** | Infants | 36 | 2-24 months | 83% (n=30) | ND | 36% (n=13) |
|  | Children | 23 | 24-60 months | 78% (n=18) | ND | 39% (n=9) |
|  | Adults | 91 | 18-60 years | +/- ** | ND | 51% (n=47) |

ND = Not Done. \*While QFT and TST were not performed, the infection status in Oregon is likely to be negative based on epidemiology and results of an in-house ESAT-6/CFP-10 IFN-γ ELISPOT assay. \*\*This information is not known.

### Frequencies of tetramer-defined MR1T cell and phenotypically defined MAIT cell populations increase with age irrespective of BCG vaccination

We defined MR1T cells as CD3^+^ T cells positively labeled with the MR1-5-OP-RU tetramer. The MR1-6FP tetramer was as a negative control tetramer (Figure 1A). The MR1-6-FP tetramer stained a different population of cells than the MR1-5-OP-RU tetramer (Figure S1). In addition, MAIT cells, were defined phenotypically as CD3^+^ T cells that co-expressed TRAV1-2 and high levels of CD161 (CD161^++^) and CD26 (CD26^++^) (Figure 1A and Figure S2). This phenotypic definition of MAIT cells is concordant with the vast majority of MR1-5-OP-RU tetramer-defined MR1T cells in adults (Sharma et al., 2015; Meermeier et al., 2016). We determined frequencies of MR1T cells and the phenotypically-defined MAIT cells in all cohorts (Figure 1B). Because of limited availability of the MR1-5-OP-RU tetramer, determination of MR1-5-OP-RU tetramer^+^ MR1T cell frequencies in the Ugandan cohorts was performed on a subset of 10 donors (Figure 1B and Figure S3). In all cohorts, both MR1-5-OP-RU tetramer-defined MR1T cell and phenotypically-defined MAIT cell frequencies (Figure 1B) and absolute cell numbers (Figure S4) increased with age.

**Figure 1.**
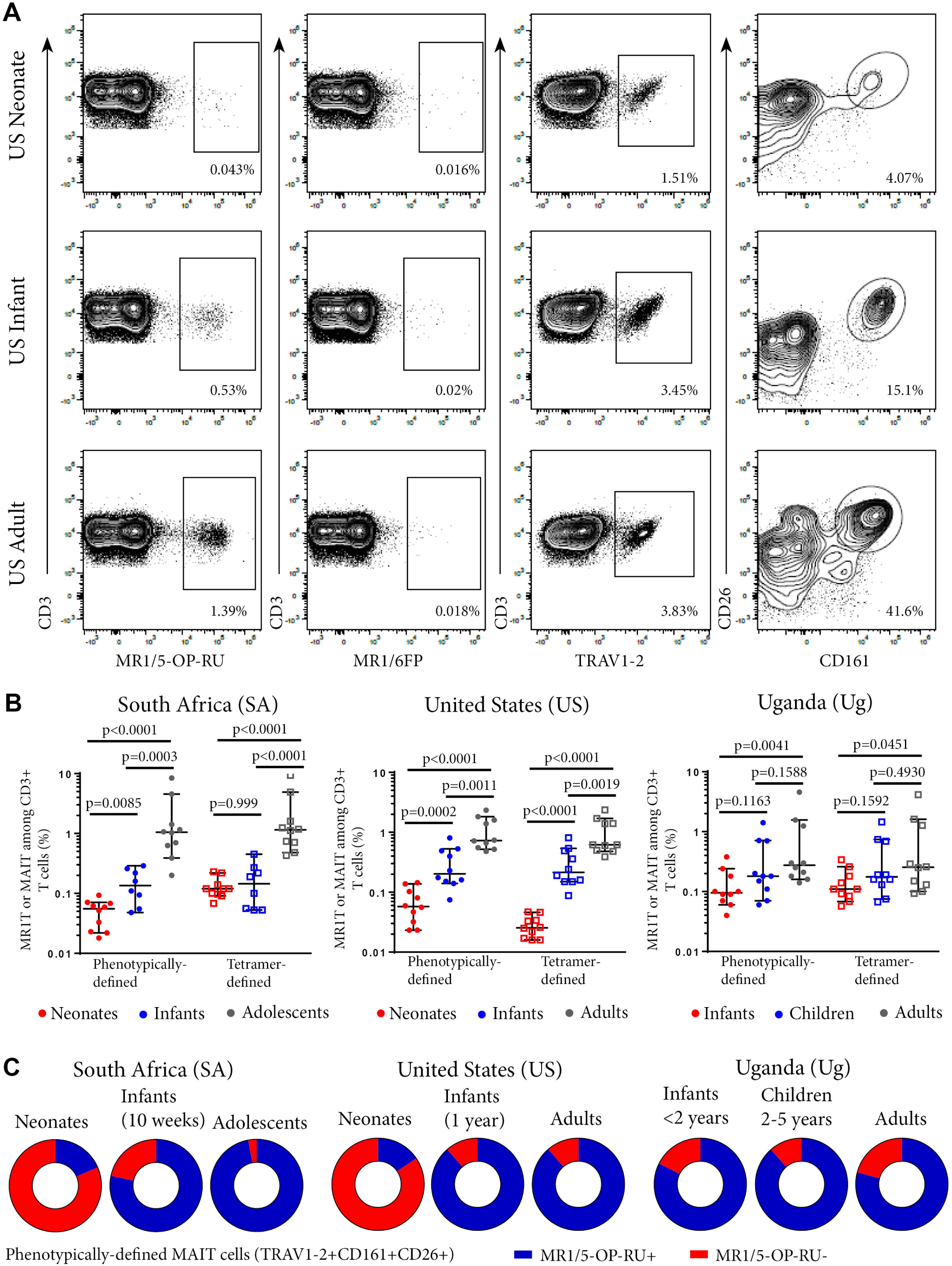
Frequencies of tetramer-defined MR1T cell and phenotypically defined MAIT cell populations in peripheral blood from individuals of different ages. PBMC or CBMC were stained with a live/dead discriminator, antibodies to CD3, CD4, CD8, TRAV1-2, CD26, CD161, and either the MR1-5-OP-RU or MR1-6FP tetramers. Live, CD3^+^ lymphocytes were gated and the frequencies of MR1-5-OP-RU^+^ or TRAV1-2^+^CD161^++^CD26^++^ cells as a percentage of CD3^+^ lymphocytes were determined (gating strategy in Figure S2). **A**. Flow cytometry plots showing CD3^+^ T cell staining with MR1-5-OP-RU tetramer, MR1-6FP tetramer, TRAV1-2 or CD26/CD161 of samples from a representative US adult, infant and neonate. **B.** Frequencies of tetramer-defined (CD3^+^MR1-5-OP-RU^+^) and phenotypically-defined (CD3^+^TRAV1-2^+^ CD161^++^CD26^++^) MAIT cells in neonates, 10 week-old infants and adolescents from South Africa, neonates, 12 month-old infants and adults from the United States and infants (0-2 years old), children (2-5 years old) and adults from Uganda (all cohorts, *n*=10). Mann-Whitney u-tests were used to test differences between groups. Horizontal lines depict the median and the error bars the 95% confidence interval. **C.** Relative proportions of median phenotypically-defined (CD3^+^TRAV1-2^+^CD161^++^CD26^++^) MAIT cells that are MR1-5-OP-RU^+^ (Blue) and MR1-5-OP-RU^−^ (Red) for each age group at each site.

### Tetramer-defined MR1T cell and phenotypically-defined MAIT cell populations are concordant in adults, adolescents and infants but discordant in neonates

We next determined the degree of concordance between tetramer-defined MR1T cell and phenotypically-defined MAIT cell populations. The majority of the phenotypically-defined MAIT cell population (TRAV1-2^+^CD161^++^CD26^++^ cells) also stained with the MR1-5-OP-RU tetramer in infants, adolescents and adults, demonstrating high concordance between tetramer-defined MR1T cells and phenotypically-defined MAIT cells in these cohorts. (Figure 1C). However, in cord blood from both SA and US neonates, these two populations were discordant as the minority of TRAV1-2^+^ CD161^++^CD26^++^ MAIT cells also stained with the MR1T tetramer (Figure 1C and Figure S5).

### The MR1-5-OP-RU tetramer^+^ subset of phenotypically-defined MAIT cells contain the functional population

To investigate whether MR1-5-OP-RU tetramer-defined MR1T cells and phenotypically-defined MAIT cells have different functional capacity, we measured TNF expression in the US cohort by intracellular cytokine staining (ICS) and flow cytometry following incubation with *M. smegmatis-*infected A549 cells. Among all phenotypically defined TRAV1-2^+^CD161^++^CD26^++^ MAIT cells, only the MR1-5-OP-RU tetramer^+^ subset expressed TNF; MR1-5-OP-RU tetramer-negative cells, by contrast, did not appear to respond to *M. smegmatis* stimulation (Figure 2). Because of this result, we focused on the functional, CD3^+^ MR1-5-OP-RU tetramer^+^ MR1T cells for subsequent experiments.

**Figure 2.**
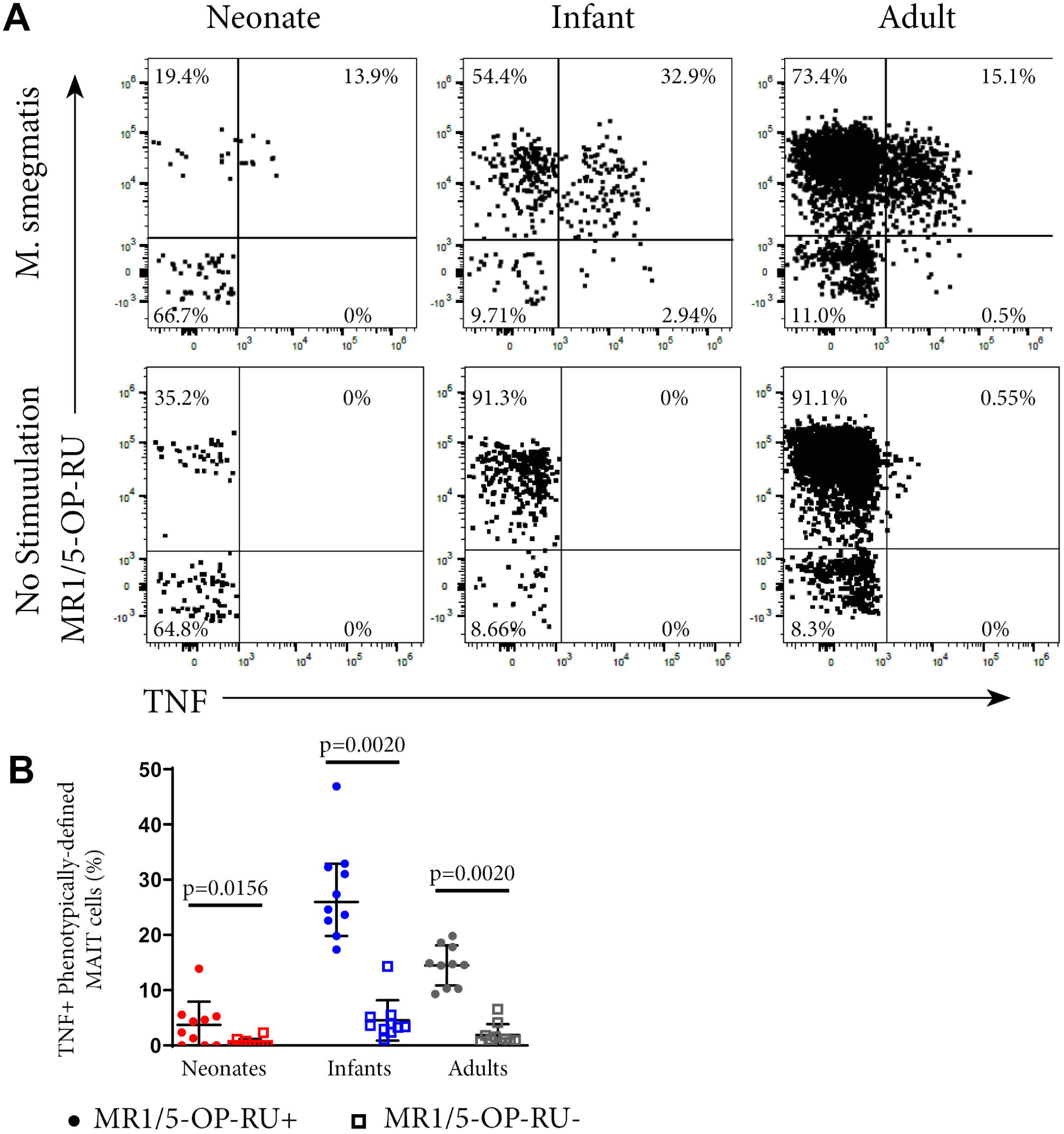
Functional analysis of MAIT cells in individuals of different ages. PBMC or CBMC from the US cohort were incubated overnight with *M. smegmatis*-infected A549 cells or uninfected A549 cells and then stained with the MR1-5-OP-RU or MR1-6FP tetramers, followed by staining with a live/dead discriminator and antibodies to TCRγδ, CD3, CD4, CD8, TRAV1-2, CD26 and CD161. ICS was then performed and the cells stained for TNF. **A.** Dot plots showing representative co-staining of live, TCRγδ^−^CD3^+^TRAV1-2^+^CD161^++^CD26^++^ cells with MR1-5-OP-RU and TNF in *M. smegmatis* stimulated or unstimulated samples. The gating strategy is in Figure S6. Examples of the TNF response in a neonate, infant and adult are shown. **B.** Frequencies of phenotypically-defined (CD3^+^TRAV1-2^+^CD161^++^CD26^++^) MAIT cells that are TNF^+^MR1-5-OP-RU^+^ or TNF^+^MR1-5-OP-RU^−^ are shown. Horizontal lines depict the median and the error bars the 95% confidence interval. Wilcoxon-rank sum was used to test differences within the same cohort.

### Neonates have higher proportions of CD4^+^ and TRAV1-2^−^ MR1T populations compared to infants, adolescents and adults

To explore the diversity of MR1T cells and how this may change during early development, we applied the dimensionality reduction method, t-distributed stochastic neighbor embedding (tSNE), on MR1-5-OP-RU tetramer^+^ MR1T cells. Five discrete phenotypic MR1T cell clusters were identified in both the SA and US cohorts (Figure 3A-D), the relative proportions of which varied with age (Figure 3B and D). Specifically, the CD4^+^ (Clusters 3 and 4) and TRAV1-2^−^CD26^−^ CD161^−^ (Clusters 1, 2 and 3) MR1T cell clusters predominated in neonates, while CD4^−^CD8^+/−^ TRAV1-2^+^CD26^+^CD161^+^ cluster (Cluster 5) was predominant in infant, adolescent, and adult cohorts from both the US and SA (Figure 3B and D). Relative proportions of MR1T cell clusters in 10 week old (SA) and 12 month old (US) infants were more similar to those in adults and adolescents than to those in neonates, consistent with the hypothesis that MR1T cells expand rapidly upon exposure to MR1T cell antigens (Figure 3E and F). It was also noteworthy that CD4^−^ CD8^+/−^TRAV1-2^+^CD161^+^CD26^+^ MR1T cells that fell into cluster 5 comprised a much lower percentage of the MR1Ts in cord blood, while forming the dominant phenotype in 12-month-old (US) infants. In the 10 week old (SA) infants, this cluster had started to appear but was not yet “fully developed”, suggesting that the transition from a predominantly CD4^+^ and TRAV1-2^−^ population in neonates, to a predominantly TRAV1-2^+^CD161^++^CD26^++^ CD8^+^ population occurs during the first months of life. We also note that in the US cohort only there is a population that is positive for all markers (Cluster 6) that is present at low frequency in all ages.

**Figure 3.**
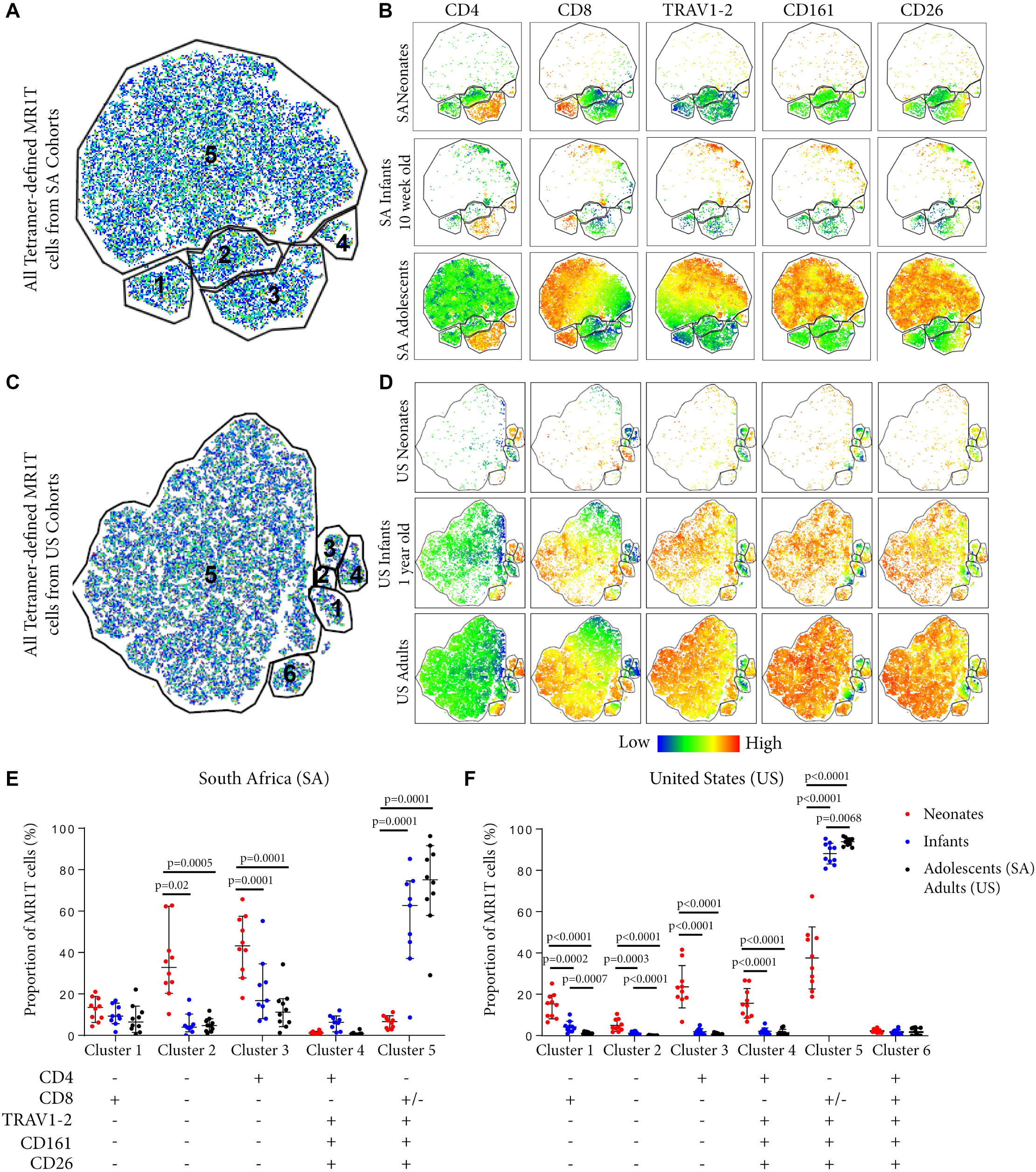
Phenotypic diversity of MR1T cells in individuals of different ages. t-Distributed stochastic neighbor embedding (tSNE) plots showing MR1-5-OP-RU tetramer^+^CD3^+^ T cells (gating strategy in Figure S2) from the South African (SA) or United States (US) cohorts (Table1). Plots represent cells from 10 samples from each age group. **A.** and **C.** The five clusters identified in both sites are shown in **A** (SA) and **C** (US). **B.** and **D.** Dot plots colored by the relative expression level of each marker (CD4, CD8, TRAV1-2, CD161 or CD26) in tetramer-defined MR1T cells for each age group in **B** (SA) and **D** (US). Red = highest expression, blue = lowest expression. **E.** and **F.** The proportion of each cluster of tetramer-defined MR1T cells by age group is shown in **E** (SA) and **F** (US). Horizontal lines depict the median and the error bars the 95% confidence interval. Mann-Whitney u tests were used to test differences between groups.

### MR1T cells become more functional with age

Next, we explored how tetramer-defined MR1T cells developed functionally with age in the US cohort by tracking the TNF-expressing population following *M. smegmatis* or PMA/Ionomycin stimulation. TNF-expressing tetramer^+^ MR1T cells in neonates were present at very low frequencies (Figure 2B) and represented a much smaller proportion of the total tetramer^+^ MR1T cell population than in infants and adults (Figure 4A and B). Proportions of tetramer^+^ MR1T cells responding to either mycobacteria or PMA/Ionomycin were higher in infants than adults (p=0.0007 and p=0.0021, respectively) or neonates (p<0.0001 for both stimulations). However, because tetramer^+^ MR1T cell frequencies were higher in adults than infants (Figure 1B), overall frequencies of TNF-producing MR1T cells were highest in adults. To explore the functional capacity of MR1T cells relative to other T cell subsets within each cohort, we compared proportions of TNF-expressing MR1T cells (CD3^+^TCRγδ^−^MR1-5-OP-RU^+^) with TCRγδ T cells (CD3^+^TCRγδ^+^ MR1-5OPRU^−^), CD4^+^ T cells (CD3^+^TCRγδ^−^MR1-5OPRU^−^CD4^+^) and CD8^+^ T cells (CD3^+^TCRγδ^−^MR1-5OPRU^−^CD4^+^) after PMA/Ionomycin stimulation (Figure 4C). In neonates, proportions of TNF-expressing MR1T cells were significantly higher than proportions of TCRγδ T cells and CD8^+^ T cells. In infants and adults, proportions of TNF-expressing MR1T cells were significantly higher than TCRγδ^+^, CD4^+^ and CD8^+^ T cells (Figure 4C).

**Figure 4.**
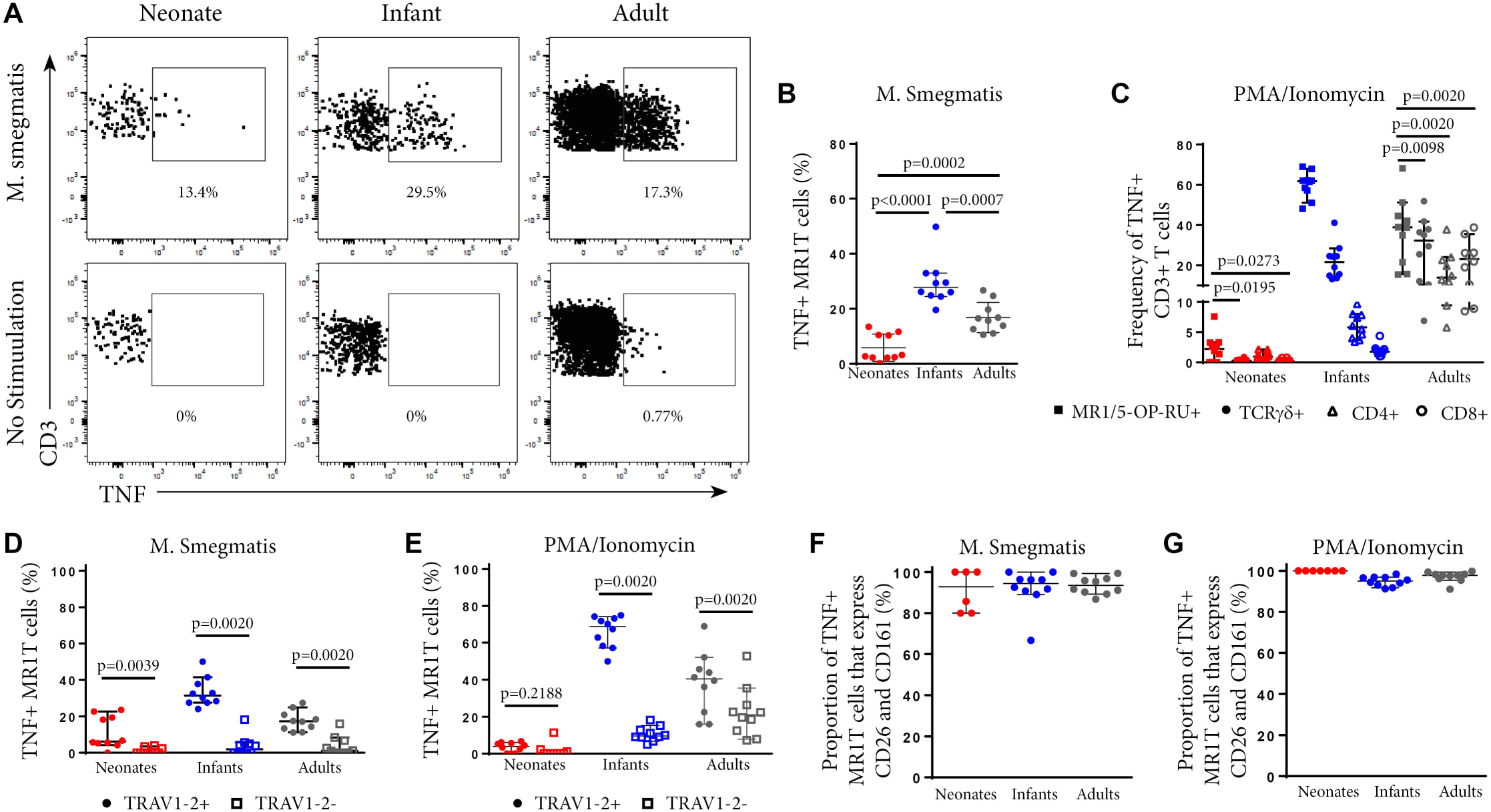
Phenotypic analysis of functional MR1T cells at different ages. PBMC or CBMC from the US cohort were incubated overnight with *M. smegmatis*-infected A549 cells, uninfected A549 cells, or uninfected A549 cells and PMA/ionomycin. All cells were then stained with the MR1-5-OP-RU or MR1-6FP tetramers, followed by a live/dead discriminator and antibodies to TCRγδ, CD3, CD4, CD8, TRAV1-2, CD26 and CD161. ICS was then performed and the cells stained for TNF. Live, TCRγδ^−^CD3^+^MR1-5-OP-RU^+^ cells were gated and the TNF^+^ percentage determined in both the *M. smegmatis* stimulated and unstimulated conditions. The gating strategy is shown in Figure S6. **A.** Dot plots showing TNF expression by live, TCRγδ^−^MR1-5-OP-RU^+^ CD3^+^ cells in the same representative neonate, infant and adult as Figure 2A. **B.** Background subtracted frequencies of TNF^+^ MR1-5-OP-RU^+^ CD3^+^ cells in response to *M. smegmatis*-infected A549 cells as a percentage of total MR1-5-OP-RU^+^ CD3^+^ cells are shown. Mann-Whitney u-tests were used to test differences between groups. **C.** Background subtracted frequencies of TNF^+^ cells to PMA/ionomycin among different T cell subpopulations, 1) MR1T cells (CD3^+^TCRγδ^−^ MR1-5-OP-RU^+^ cells); 2) γδ T cells (CD3^+^TCRγδ^+^MR1-5-OP-RU^−^ cells); 3) CD8^+^ T cells (CD3^+^TCRγδ^−^MR1-5-OP-RU^−^CD8^+^ cells); and 4) CD4^+^ T cells (CD3^+^TCRγδ^−^MR1-5-OP-RU^−^ CD4^+^ cells). Wilcoxon-rank sum was used to test differences within the same cohort. **D.** and **E.** Frequencies of TNF^+^TRAV1-2^+^ MR1-5-OP-RU^+^ CD3^+^ and TNF^+^TRAV1-2^−^ MR1-5-OP-RU^+^ CD3^+^ cells in response to *M. smegmatis* infected A549 cells (**D**) or PMA/ionomycin stimulation (**E**) minus the background frequencies of TNF^+^TRAV1-2^+^/- MR1-5-OP-RU^+^ CD3^+^ cells. Wilcoxon-rank sum was used to test differences within the same cohort. **F.** and **G.** Frequencies of CD161^++^CD26^++^ cells of the TNF^+^TRAV1-2^+^MR1-5-OP-RU^+^CD3^+^ cells in response to *M. smegmatis* infected A549 cells (**F**) or PMA/ionomycin stimulation (**G**) are shown. The median and 95% confidence intervals are shown in each graph. When the 95% confidence intervals do not overlap between conditions, those conditions are considered statistically significant.

### Functional CD3^+^MR1-5-OP-RU^+^ MR1T cells are TRAV1-2^+^

Next, we investigated TRAV1-2 expression among functional, TNF-expressing CD3^+^MR1-5-OP-RU^+^ MR1T cells. Only TRAV1-2^+^ MR1T cells produced TNF in response to mycobacterial stimulation while TRAV1-2^−^ MR1T cells were not functional (Figure 4D). This pattern was unique to mycobacterial stimulation since TNF-expressing TRAV1-2^−^ MR1T cells were observed in response to PMA/Ionomycin stimulation, particularly in infants and adults (Figure 4E). Interestingly, the vast majority of these functional, TNF-expressing TRAV1-2^+^ MR1T cells also highly expressed the phenotypic markers, CD26 and CD161 (Figure 4F-G). Thus, the CD3^+^MR1-5-OP-RU^+^TRAV1-2^+^CD161^++^CD26^++^ MR1T cell population contains the functional MR1T cells that respond to mycobacteria. The frequency of this population increases very quickly with age.

### TRAV1-2^+^ MR1T cells lose naïve markers more rapidly with age than TRAV1-2^−^ MR1T cells or non-MR1T cells

To characterize the development of MR1T cells further, we also measured expression of CD27 and CD161, markers previously described to indicate MR1T cell maturation (Koay et al., 2018), and CD45RA, CCR7, CD38 and CD56 on tetramer-defined MR1Ts (Figures 5-6). A population of immature MR1T cells, defined as CD27^+^CD161^−^ cells, was observed in neonates but not in infants and adults, as previously described (Koay et al., 2016). This was also reflected by expression of the more traditional T cell maturation markers, CD45RA, CCR7 and CD38 (Figure 6 A, C, E). Most tetramer-defined MR1T cells in neonates displayed a CD45RA^+^CCR7^+^CD38^+^ “naïve” phenotype, but this phenotype was lost more rapidly with age on TRAV1-2^+^ MR1T cells than on TRAV1-2^−^ MR1T cells or non-MR1T cells (Figure 6A-F). The inverse was true for the activation marker, CD56, which was increased in neonatal, infant and adult TRAV1-2^+^ tetramer-defined MR1T cells compared to TRAV1-2^−^ tetramer-defined MR1T cells and non-MR1T cells (Figure 6G-H). These results are consistent with our finding that TRAV1-2^+^ tetramer-defined MR1T cells display functional capacity in all age groups, even though they represent the minority of neonatal MR1T cells.

**Figure 5.**
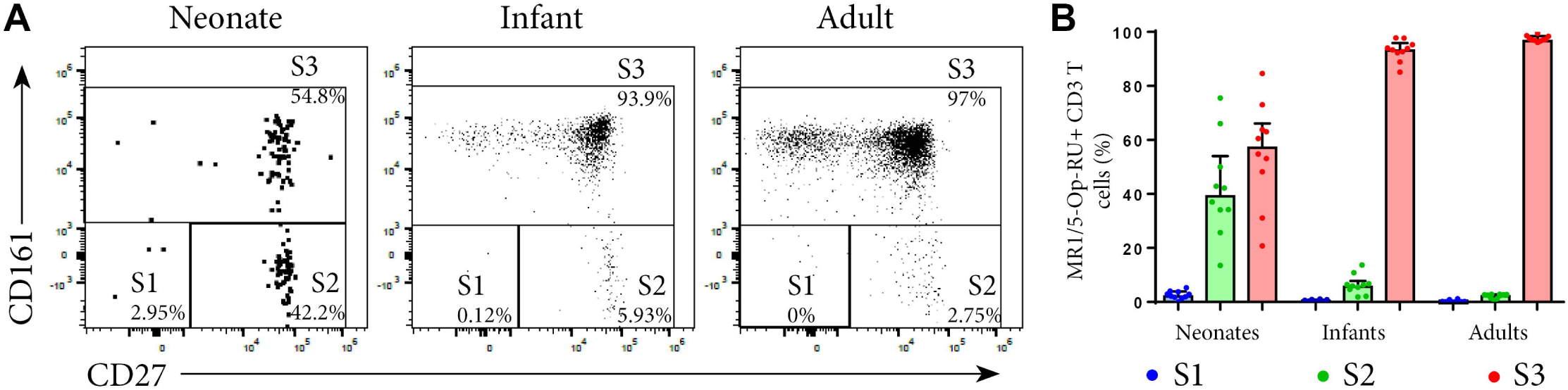
Analysis of MR1T cell development by measuring maturation markers. PBMC or CBMC from the US were stained as in Figure 1 with the addition of an antibody to CD27. Live, CD3^+^MR1-5-OP-RU^+^ cells (gating strategy in Figure S2) were gated for the maturation pattern of S1, S2 and S3 described in (Koay et al., 2016). S1 = CD161^−^CD27^−^, S2=CD161^−^CD27^+^, S3=CD161^+^CD27^+/−^. **A.** Gating of MR1T cells for S1, S2 and S3 in a representative neonate, infant and adult. **B.** Frequencies of S1, S2, and S3 MR1T cells as a percentage of CD3^+^ MR1-5-OP-RU^+^ cells are shown for U.S. neonates, infants and adults. Bars represent the median and the error bars the 95% confidence interval.

**Figure 6.**
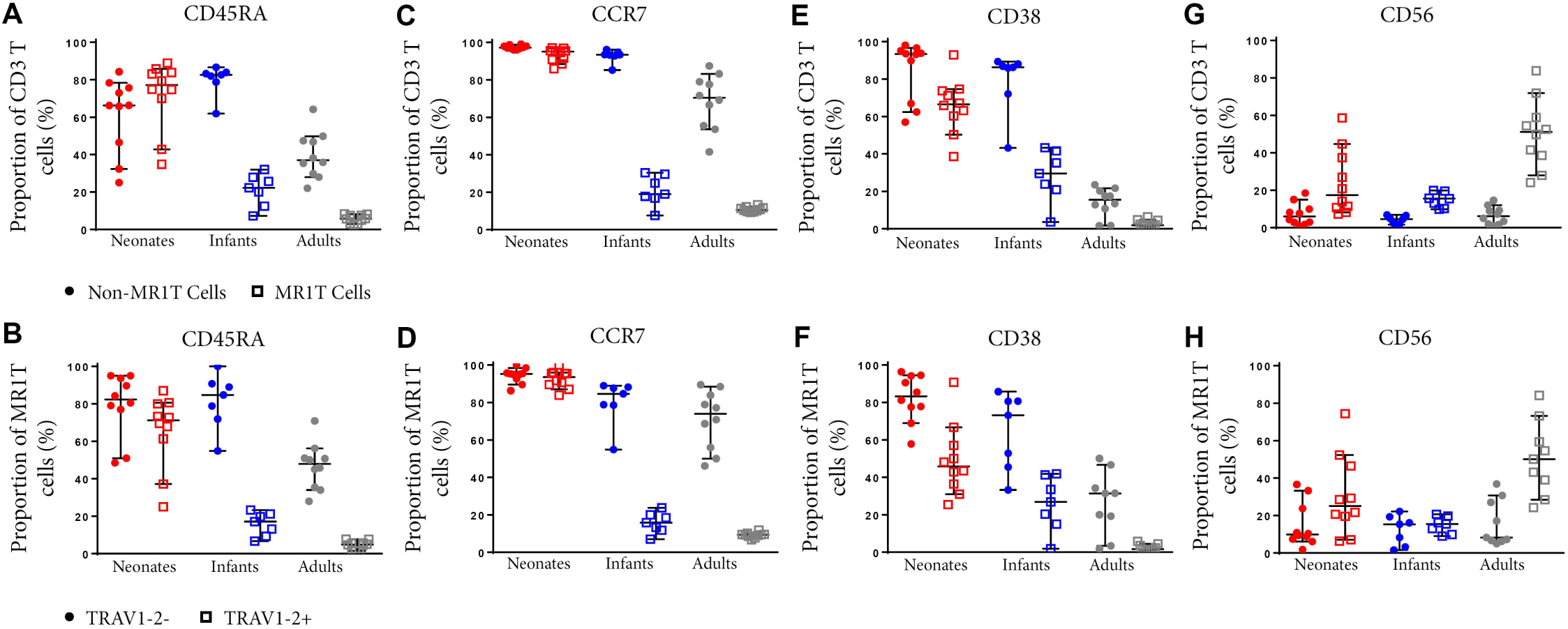
Comparative phenotypic analysis of MR1T and non-MR1T cells or TRAV1-2^+^ and TRAV1-2^−^ MR1T cells at different ages. PBMC or CBMC from the US were stained as in Figure 1 with the addition of antibodies to either CD45RA (**A, B**), CCR7 (**C, D**), CD38 (**E, F**) or CD56 (**G, H**). The gating strategy is shown in Figure S2 and examples of the staining of each marker is shown in Figure S7. Frequencies of cells expressing each marker in non-MR1T cells (CD3^+^MR1-5-OP-RU^−^) or MR1T cells (CD3^+^MR1-5-OP-RU^+^) in each age group (**A, C, E, G**). The frequency of each marker expressed in TRAV1-2^+^ or TRAV1-2^−^ MR1T cells is shown in each age group (**B, D, F, H**). The median and 95% confidence intervals are shown in each graph. When the 95% confidence intervals do not overlap between conditions, those conditions are considered statistically significant.

## Discussion

We characterized the development of MR1T cells during early childhood in three cohorts, two from Africa and one in the USA, to elucidate phenotypic and functional changes that occur after birth and during the first few years of life. A better understanding of MR1T cell development is critical given the likely role of MR1T cells as innate-like effectors that act as key mediators of immunity in newborns, who are at very high risk of infection, including *M. tuberculosis.* We found that neonates had relatively low frequencies of MR1T cells with high phenotypic diversity, which included considerable proportions of CD4^+^ and TRAV1-2^−^ cells. These patterns changed as early as 10 weeks after birth, and became more prominent at 1-2 years of age, adolescence and ultimately by adulthood. This included higher frequencies of MR1T cells in peripheral blood, characterized by a preferential expansion in TRAV1-2^+^CD8^+^ MR1T cells such that TRAV1-2^+^CD8^+^MR1T cells constituted the vast majority of the tetramer^+^ MR1T cell population. The development also included changes in expression of maturation markers and MR1T cell effector function, since MR1T cells from older individuals had lower expression of the naïve T cell markers, CD45RA, CCR7 and CD38, and an increased ability to express TNF.

Our finding that MR1T cells are found at low frequency in cord blood and increase in frequency with age confirms our previous observations, as well as those of others, who have utilized either the MR1-5-OP-RU tetramer or phenotypic characterization of MAIT cells to demonstrate the expansion of these cell populations from birth through adulthood (Martin et al., 2009; Walker et al., 2012; Gold et al., 2013; Walker et al., 2014; Koay et al., 2016; Ben Youssef et al., 2018; Gherardin et al., 2018). The very rapid changes in MR1T cells seen after birth are striking and were clearly seen in both African and American settings. This finding rules out the possibility that BCG substantially contributes to these changes, since BCG is not routinely administered at birth in the USA and is consistent with the lack of differences in MAIT cell magnitude or function between young infants either BCG immunized or unimmunized (Suliman et al., 2019). Rather, it is more likely that changes occur upon the transition from the limited microbial exposure in the womb to the postnatal environment, accompanied by the rapid colonization of the gut microbiome. MR1T cells are absent in germ-free mice and depend on the colonization of the gut microbiome with normal intestinal flora, which produce riboflavin (Le Bourhis et al., 2013; Harriff et al., 2018; Legoux et al., 2019). We suggest that this colonization of the gut microbiome is the driving force that accounts for the increase and phenotypic changes in MR1T cells in blood after birth and the gain in functional capacity, though environmental exposure and infection could also contribute.

While we confirm our previous work that phenotypic definition of MAIT cells accurately identifies functional MR1T cells in adults, we find that there is a high level of discordance in neonates in part due to MR1-5-OP-RU tetramer^+^ cells that do not express TRAV1-2, and/or high levels of CD161 or CD26. These cells are by definition MR1-restricted T cells as they bind the MR1-5-OP-RU tetramer. However, these cells are poorly reactive to mycobacteria, suggesting that they either recognize a different antigen or represent functionally immature MR1T cells. The poor PMA/Ionomycin-induced TNF secretion of these cells supports the latter. However, because our only measure of MR1T cell responses was TNF production, we may have failed to detect other responses characterized by the production of alternative cytokines or cytolytic markers.

We also show that neonates possess MR1-5-OP-RU tetramer^−^ cells that phenotypically resemble MAIT cells (the discordant MR1-5-OP-RU tetramer^−^ phenotypically defined TRAV1-2^+^CD161^++^CD26^++^ MAIT cell population). As Fergusson *et al* suggested that distinct T subsets that express CD161 share a common transcriptional profile characteristic of typical MR1T cells (Fergusson et al., 2014), it is certainly possible that these CD161^++^ cells are MR1-restricted or share common attributes with MR1T cells. Further characterization of this neonatal T cell population is required to determine whether or not these cells are indeed MR1-restricted and if so, if these cells are immature or recognize distinct microbes, or produce a different cytokine not detected in our functional assay. Given that, in the mouse model, 5-OP-RU produced by the microbiome is necessary and sufficient to drive the expansion of typical MR1T cells (Legoux et al., 2019), it is perhaps expected that with age tetramer-positive MR1T TRAV1-2^+^ cells would quickly become the predominant MR1T cell population, while tetramer negative MR1T cells might persist as a minority population of naïve T cells.

Consistent with our prior findings (Gold et al., 2013) and those of others (Walker et al., 2012; Ben Youssef et al., 2018), our phenotypic analysis of MR1-5-OP-RU tetramer^+^ cells in neonates demonstrates that MR1T cells expressed a naïve cell surface phenotype (CD45RA^+^CCR7^+^CD38^+^) and included a prominent subset of CD4^+^ cells. It is noteworthy that these MR1T cells rapidly lost expression of CD45RA, CCR7 and CD38 and acquired a phenotype consistent with those observed in antigen-experienced memory T cells much more rapidly than other types of T cells (Chua et al., 2012; Tsukamoto et al., 2013; Lindestam Arlehamn et al., 2014).

While Koay *et al* showed the functional potential of neonatal MR1T cells that produced TNF following stimulation with PMA/Ionomycin, we show for the first time the presence of functional microbe-reactive MR1T cells in neonates, albeit at low frequency. Virtually all of the mycobacteria-reactive TNF-producing MR1T cells were contained within the TRAV1-2^+^CD161^++^CD26^++^ subset of 5-OP-RU tetramer^+^ cells, which are also concordant with “Stage 3” MR1T cells defined by Koay *et al*. Although the immature CD4^+^, CD4-CD8- and/or TRAV1-2-MR1T cells were maintained through to adulthood, we observed that CD8^+^TRAV1-2^+^ MR1T cells expanded preferentially over TRAV1-2^−^ MR1T cells. This expansion coincided with the acquisition of high expression of CD26 and CD161 as markers of mature functional MR1 T cells. Of note, studies evaluating the development of CD8^+^CD161^++^ T cells in humans reported that a lower proportion of this cell population expressed TRAV1-2 in neonates as compared to 2 year-olds (Fergusson et al., 2014) and adults (Walker et al., 2012; Fergusson et al., 2014), and that in adults, CD8^+^CD161^++^ T cells producing IFN-γ in response to *E. coli* expressed TRAV1-2 (Fergusson et al., 2014). While these studies did not use the MR1 tetramer, these findings are consistent with our results to the extent that the total CD8^+^ CD161^++^ population reflected MR1T cells. In addition, we show that the frequency of these functional mycobacterial-reactive MR1T cells increases ∼10-fold in the first year of life. A surprising finding was that given a comparable stimulus (PMA/Ionomycin and *M. smegmatis*), the infant MR1T cells exhibited a higher functional capacity than the adults. Interestingly, we demonstrated that relative to other T cell subsets, MR1T cells had the highest capacity to secrete the pro-inflammatory cytokine, TNF, in infants. Thus, MR1T cells may represent an important source of pro-inflammatory effectors in infants, who are more vulnerable to serious bacterial infections in the first two years of life, relative to older children and adults.

Regarding newborns, it is not known to what degree the naïve phenotype and low functionality of MR1T cells contributes to the very high susceptibility to bacterial infection in the first two months of life (Paul K Sue, 2018). Since MR1T cells recognize important childhood bacterial pathogens, including *S. aureus, S. pneumoniae and enteric pathogens (e.g. E. coli, Klebsiella, Salmonella species*, it is plausible that the low frequency of functional MR1T cells during the neonatal period contributes to neonatal susceptibility to infection. At the very least, our findings add to the well-described differences in adaptive and innate immunity between neonates and adults (Paul K Sue, 2018).

With regard to *M. tuberculosis*, there is mounting evidence that supports an important host defense role for MR1T cells, especially during early infection. Upon encounter with mycobacterial-infected cells, MR1T cells produce pro-inflammatory cytokines such as IFN-γ and TNF (Gold et al., 2010), known to be essential mediators to host defense against *M. tuberculosis* (Lindestam Arlehamn et al., 2014). MR1-deficient mice control BCG less well than wild type mice (Chua et al., 2012). In humans, MR1 polymorphisms are associated with susceptibility to TB meningitis (Seshadri et al., 2017). It is tempting to speculate, in light of the high susceptibility of infants to TB, and the phenotypic and functional attributes of MR1T cells we observed, that MR1T cells contribute to control of early infection with *M. tuberculosis* through their early and robust anti-mycobacterial response, particularly in the lung.

## Methods

### Study participants

#### University of Cape Town, Western Cape, South Africa

All participants were enrolled either at the South African Tuberculosis Vaccine Initiative Field Site, Well-Baby Clinics or Maternity Wards in the Worcester region, near Cape Town, South Africa. Pregnant mothers were enrolled for collection of cord blood immediately after childbirth by Cesarean section. Mothers who may have been exposed to *M. tuberculosis*, HIV infected, had any other acute or chronic disease, and receiving medication other than anesthetic given during delivery were excluded. Infants born to HIV negative mothers, and who had received BCG at birth, had no perinatal complications, or household contact with any person with TB, and did not have any acute or chronic disease were enrolled for collection of blood. Healthy adolescents of 12-18 years of age without evidence of latent Mtb infection as determined by positivity negativity for the Tuberculin Skin Test (<10mm) or QuantiFERON (<0.35IU/mL) were enrolled for blood collection. Heparinized blood was collected for cord blood mononuclear cell (CBMC) and peripheral blood mononuclear cells (PBMC) isolation by means of density gradient centrifugation.

#### Oregon Health and Science University, Portland, OR, USA

Umbilical cord blood was obtained from the placenta of uncomplicated term pregnancies after delivery into citrate CPT tubes (BD). Blood was processed per manufacturer’s instructions within 24hr of collection, and resulting CBMC cryopreserved. Infants aged 1-24 months undergoing elective surgery at OHSU Doernbecher Children’s Hospital were enrolled. Infants had not received BCG vaccination, which is not part of the childhood immunization series in the United States. Blood was drawn into citrate CPT tubes (BD) and processed per manufacturer’s instructions to isolate PBMC within 24hr of collection, and PBMC subsequently cryopreserved. Adult PBMC were obtained by leukapheresis from healthy adult donors and cryopreserved. Adult participants lacked any detectable responses to *Mycobacterium tuberculosis* (Mtb) immunodominant antigens CFP-10 and ESAT-6 and therefore were assumed to be Mtb uninfected.

#### Makerere University, Kampala, Uganda

Children from age 2-60 months hospitalized with lower respiratory infection (LRTI) from etiologies other than TB were recruited from Mulago Hospital Paediatric Wards. All children were HIV-negative and had no known recent TB contacts. Children were reassessed two months following enrollment to establish vital status and ensure clinical improvement. Following completion of the study, data from all participating children was reviewed by study investigators (DAL, CLL, SK) to confirm diagnosis of non-TB LRTI. Blood was drawn into citrate CPT tubes (BD) and processed per manufacturer’s instructions to isolate PBMC within 24hr of collection, and PBMC subsequently cryopreserved. Adult participants were recruited from the National TB Treatment Center at Mulago Hospital, hospital staff, and the community surrounding Kampala, Uganda, between 2001 and 2014. Participants were HIV uninfected and included individuals with (tuberculin skin test, TST ≥ 10) or without (TST < 10 mm) evidence of Mtb infection. All had the absence of signs or symptoms of TB (prolonged cough, hemoptysis, fever, weight loss, and night sweats). Heparinized blood was collected for PBMC isolation by means of density gradient centrifugation.

#### Study approvals

The studies of all participants were conducted in accordance with the World’s Medical Association’s Declaration of Helsinki guidelines. The studies from the Western Cape, South Africa were approved by the Human Research Ethics Committee (HREC) of the University of Cape Town. Written informed consent was obtained from parents or legal guardians of adolescents and infants. Adolescents also provided written informed assent. The Institutional Review Board of Oregon Health & Science University (Portland, Oregon) approved this study of participants from Portland, OR. Written, informed consent was obtained from all adult participants and from a parent or legal guardian of infants before enrollment. For participants from Kampala, Uganda, the National Council for Science and Technology, HIV/AIDS Research Committee in Uganda, the Institutional Review Board of Oregon Health & Sciences University (Portland, OR) and the Institutional Review Board of Makerere University (Kampala, Uganda) approved this study. Written, informed consent was obtained from all adult participants and from a parent or legal guardian of infants and children before enrollment.

#### Infection of antigen presenting cells

A549 cells (ATCC CCL-185) were used as stimulators for direct ex vivo determination of mycobacterial-reactive T cells. *Mycobacterium smegmatis* (strain mc^2^155, ATCC) was grown to OD600 of 0.5 and frozen down. This was functionally titered in this assay to find the optimal amount of *M. smegmatis* (4.13μl/ml) that elicited a strong response in an adult control donor but did not reduce the viability of the antigen presenting cells. A549 cells were seeded in T25 flasks at 1.25e6 in 7.5ml of antibiotic free media. After allowing cells to adhere for 3 hours, the flasks were infected with 31μl of *M. smegmatis*. After 18 h, the cells were harvested and washed twice in RPMI/10% human serum containing antibiotic (gentamicin) before being resuspended in RPMI/10% human serum containing antibiotic (gentamicin).

#### Flow cytometry assays

To perform ICS, cryopreserved PBMC were thawed in the presence of DNAse, resuspended, in 10% heat inactivated human serum with RPMI (Lonza) at a concentration of 2 × 10^7/ml. PBMC (2 × 10^6^ cells/well) were stimulated with anti-CD28 (1 μg/ml) and anti-CD49d (1 μg/ml) (FastImmune, BD Biosciences) and 1e5 *M. smegmatis*-infected or uninfected A549 cells in RPMI/10% Human Serum for 18 h at 37°C with 5% C0_2_. During the last 12 h of this incubation, Brefeldin-A (5 μg/ml) was added. PBMC incubated with uninfected A549 cells were used to assess for background cytokine production. To determine the total functional capacity of T cells, PBMC were incubated with uninfected A549 cells and PMA/Ionomycin (PMA, 20ng/ml, Sigma; Ionomycin, 1μM, Sigma) for 3 hr. PBMC were then harvested and stained with MR1-5-OP-RU or MR1-6FP tetramers (Corbett et al., 2014) (NIH Tetramer Core) at a 1:500 dilution for 45 minutes at room temperature. All tetramers used were conjugated to R-phycoerythrin (PE) or Brilliant Violet (BV) 421. After 45 minutes, an antibody cocktail containing LIVE/DEAD Fixable Dead Cell Stain Kit (Thermo Fisher), and the surface antibodies listed in S1 Table were added for 30 minutes at 4°C. After washing, cells were fixed and permeabilized using Cytofix/Cytoperm (BD Biosciences) per manufacturer’s instructions. Cells were then stained for CD3 and TNF, then fixed in 1% PFA. Acquisition was performed using a (Beckman Coulter) Cytoflex flow cytometer with CytExpert software (BC). All flow cytometry data was analyzed using FlowJo software (TreeStar) and Prism (Graphpad).

To perform cell surface staining for phenotyping MR1T cells, cryopreserved PBMC were thawed as described above, and 1-2 × 10^6^ PBMC were stained with MR1/5-OP-RU or MR1/6FP tetramers at 1:500 for 45 minutes at room temperature. Cells were then subsequently stained with an antibody cocktail containing LIVE/DEAD Fixable Dead Cell Stain Kit (Thermo Fisher), and surface stained with the antibodies listed in S2-S4 Tables for 30 minutes at 4°C. Samples were washed and fixed with 1% PFA. Acquisition of the US samples was performed using a Cytoflex flow cytometer (Beckman) with CytExpert software (BC). For experiments with the Ugandan samples, acquisition was performed on a BD LSRFortessa. For experiments performed at SATVI, samples were acquired on a BD LSR-II flow cytometer configured with 4 lasers: solid state blue (488nm; 100mW; 3 detectors), solid state violet (405nm; 25mW; 8 detectors), HeNe gas red (635nm; 70mW; 3 detectors), and diode-pumped Coherent Compass yellow-green (532nm; 150mW; 8 detectors). Flow cytometric analysis was performed using FlowJo version 10.5.3 (Treestar, Ashland, OR, USA).

#### Data analysis and Statistics

Adjusted frequency (raw frequency of cytokine production following stimulation minus the frequency of cytokine production in resting/unstimulated condition), was used to account for the potential presence of background, or non-specific, responses. A positive cytokine response was defined as the detection of at least 0.05% cytokine-positive T cells (following background subtraction). Prism version 7 (GraphPad, La Jolla, CA, USA) was used for data analysis. A p-value < 0.05 was considered as statistically significant.

To characterize phenotypic diversity within MR1T cell populations at the different ages we performed t-Stochastic Neighbor Embedding (tSNE) on MR1-5-OP-RU tetramer^+^ defined CD3^+^ T cells. Data from MR1-5-OP-RU tetramer^+^ cells from all individuals across the age groups were concatenated into a single FCS file to perform tSNE analysis in FlowJo. Equal event sampling was not performed due to low events per individual in the cord blood and infant samples. Clustering analysis was performed based on the expression of five surface markers (CD4, CD8, TRAV1-2, CD26, and CD161). Parameter settings were fixed at 2000 iterations with a learning rate (Eta) of 10 and perplexity 30, respectively (van der Maaten and Hinton, 2008).

## Author Contributions

GMS, AG, DML, TJS and DAL contributed to the conception and/or design of the work. GMS and AG are joint first authors. TJS and DAL jointly supervised the work. MN, HMK, SK, JK, CL, EN, MS, WAH, MH and TJS contributed to clinical activities. EN, WAH, MH, DML, DAL and TJS raised grants to fund the research. GMS, AG, MEC, MDN, RBD, DML, TJS and DAL substantially contributed to the acquisition, analysis or interpretation of data and drafting of the manuscript. All authors substantially contributed to revising and critically reviewing the manuscript for important intellectual content. All authors approved the final version of this manuscript to be published and agree to be accountable for all aspects of the work.

## Acknowledgments

We would like to thank the participants who gave time and dedication to this health research; W. Henry Boom, LaShaunda Malone and Keith Chervenak, Case Western Reserve University; and Erin Merrifield, Department of Pediatrics, OHSU, for their contributions to this study.

The MR1 tetramer technology was developed jointly by Dr. James McCluskey, Dr. Jamie Rossjohn, and Dr. David Fairlie, and the material was produced by the NIH Tetramer Core Facility as permitted to be distributed by the University of Melbourne.

This manuscript has been released as a pre-print at bioRxiv, (Swarbrick et al., 2019).

