## Supplementary Materials for "Postnatal Expansion, Maturation and Functionality of MR1T Cells in Humans"

### Supplementary Material

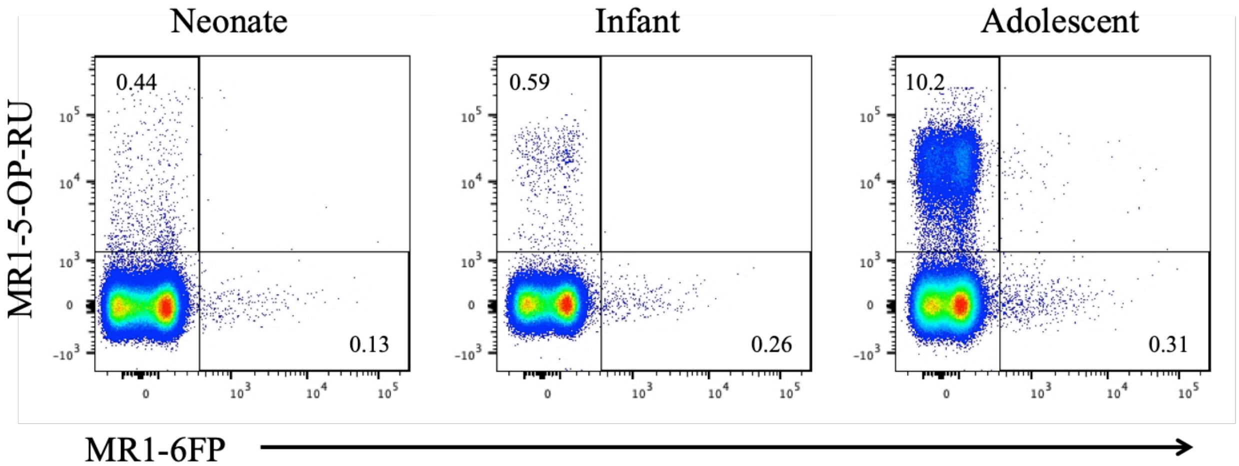


**Figure S1. Co-staining of MR1-5-OP-RU and MR1-6-FP in South African CBMC and PBMC.** Cells were stained with a live/dead discriminator, and antibodies to CD3, CD4, CD8 and the MR1-5-OP-RU and MR1-6-FP tetramers. Live, $CD3^+$ lymphocytes were gated and the staining of MR1-5-OP-RU and MR1-6-FP is shown for a representative neonate, infant and adolescent. The bimodal staining pattern observed in tetramer-negative cells in the MR1-6FP channel (tetramer was conjugated to FITC) derived from a problem with the photomultiplier tube detector. This pattern was resolved upon replacement of the detector. Importantly, this issue did not affect the flow cytometry readouts such as identification of tetramer+ cells or compensation requirements and therefore had no effect on interpretation of the results in this study.

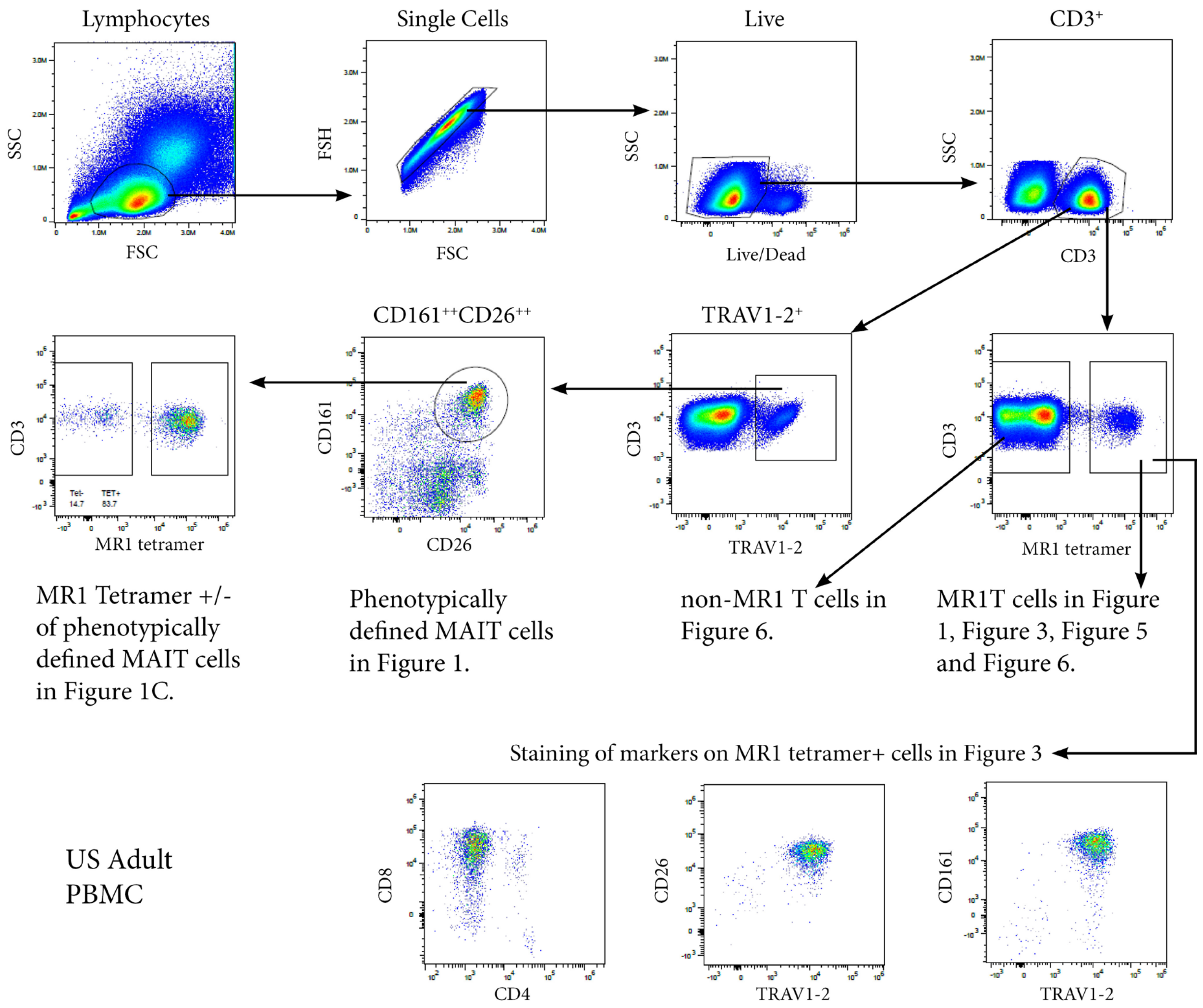


**Figure S2. Gating strategy for MR1T cells and phenotypically-defined MAIT cells.** PBMC or CBMC were stained with a live/dead discriminator, antibodies to CD3, CD4, CD8, TRAV1-2, CD26, CD161, and either the MR1-5-OP-RU or MR1-6FP tetramers. Live, $CD3^{+}$ lymphocytes were gated and the frequencies of MR1-5-OP-$RU^{+}$ or TRAV1-$2^{+}CD161^{++}CD26^{++}$ was determined. Shown in the example is PBMC from an adult from the US cohort.

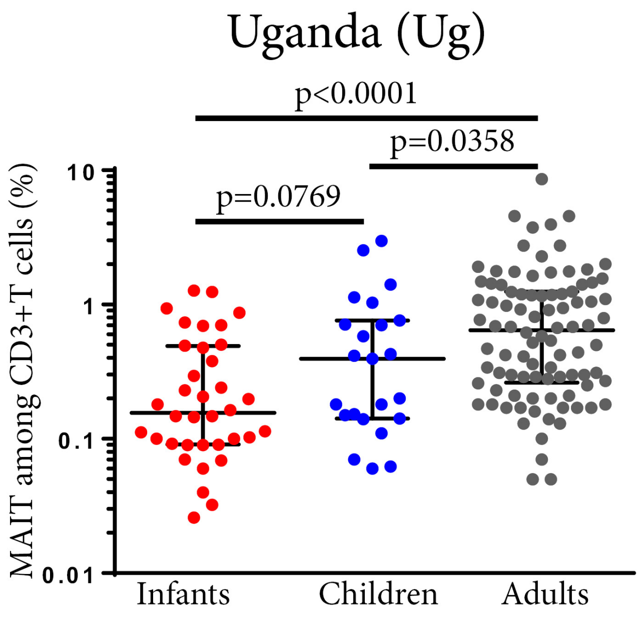


**Figure S3. Frequencies of phenotypically-defined ($CD3^{+}TRAV1\text{-}2^{+}CD161^{++}CD26^{++}$) MAIT cells in peripheral blood from the full sample size of Ugandan individuals of different ages.** PBMC were stained with a live/dead discriminator, and antibodies to CD3, CD4, CD8, TRAV1-2, CD26 and CD161. Live, $CD3^{+}$ lymphocytes were gated and the percentage of $TRAV1\text{-}2^{+}$ $CD161^{++}CD26^{++}$ cells was determined. Horizontal lines depict the median and the error bars the 95% confidence interval. Mann-Whitney u-tests were used to test differences between groups.

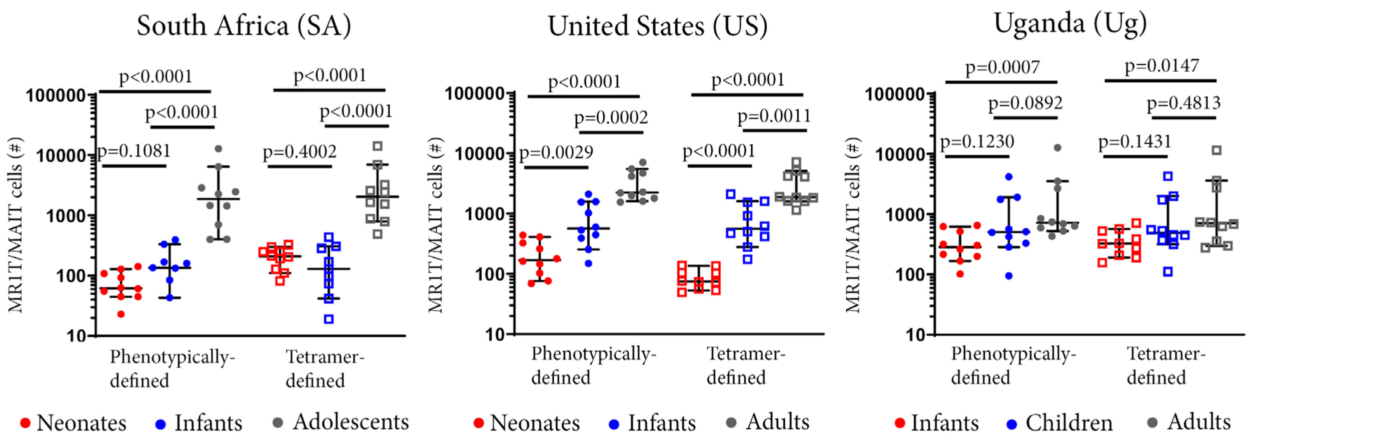


**Figure S4. Absolute numbers of CD3$^+$ tetramer-defined MR1T cell and phenotypically defined MAIT cell populations in peripheral blood from individuals of different ages.** PBMC or CBMC were stained with a live/dead discriminator, antibodies to CD3, CD4, CD8, TRAV1-2, CD26, CD161, and either the MR1-5-OP-RU or MR1-6FP tetramers. Live, CD3$^+$ lymphocytes were gated and the numbers of MR1-5-OP-RU$^+$ or TRAV1-2$^+$CD161$^{++}$CD26$^{++}$ cells were determined (gating strategy in Figure S2) in neonates, 10 week-old infants and adolescents from South Africa, neonates, 12 month-old infants and adults from the United States and infants (0-2 years old), children (2-5 years old) and adults from Uganda (all cohorts, *n*=10). Mann-Whitney u-tests were used to test differences between groups. Horizontal lines depict the median and the error bars the 95% confidence interval.

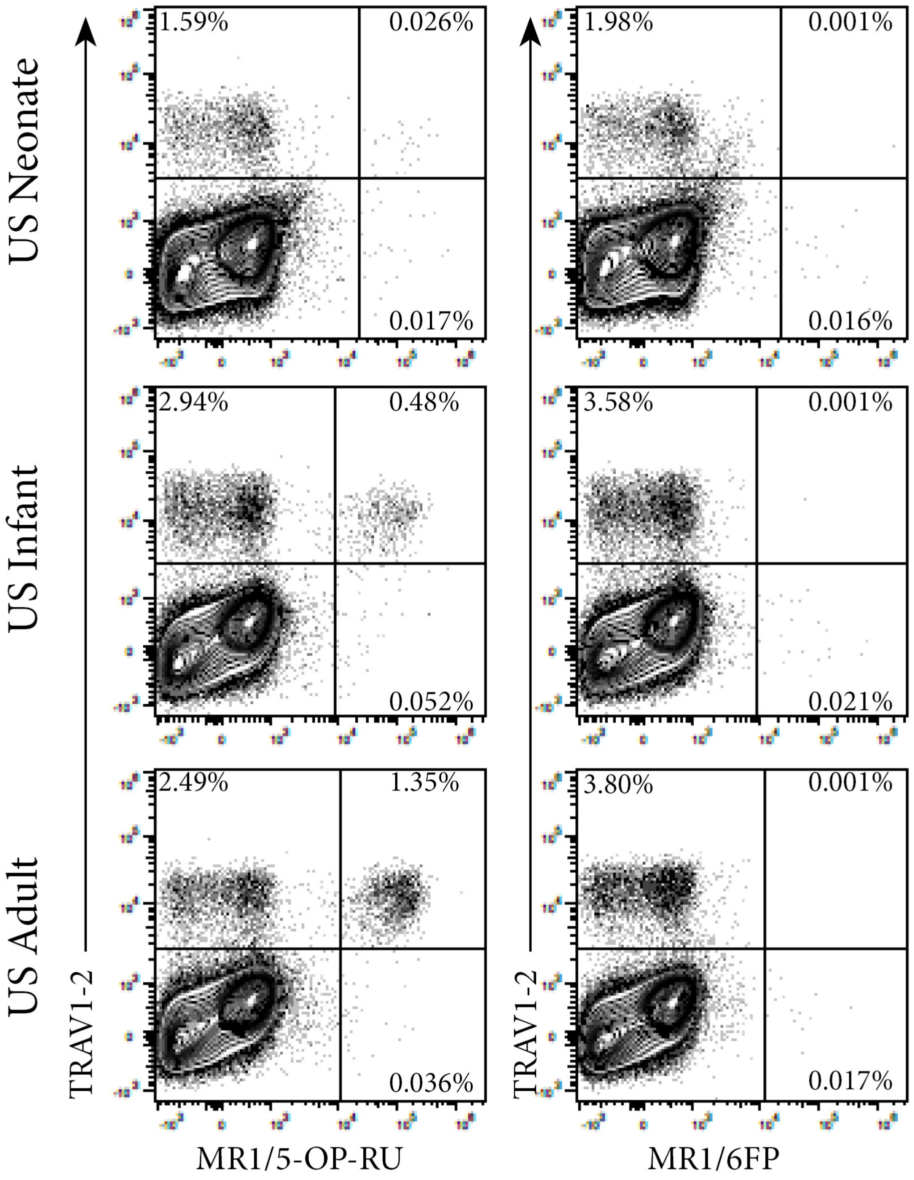


**Figure S5. Co-staining of MR1 tetramers and TRAV1-2.** PBMC or CBMC were stained with a live/dead discriminator, antibodies to CD3, CD4, CD8, TRAV1-2, CD26, CD161, and either the MR1-5-OP-RU or MR1-6FP tetramers. Live, $CD3^+$ lymphocytes were gated. Shown are flow cytometry plots showing $CD3^+$ T cell staining with MR1-5-OP-RU tetramer or MR1-6FP tetramer and TRAV1-2 of samples from the same representative US adult, infant and neonate shown in Figure 1A.

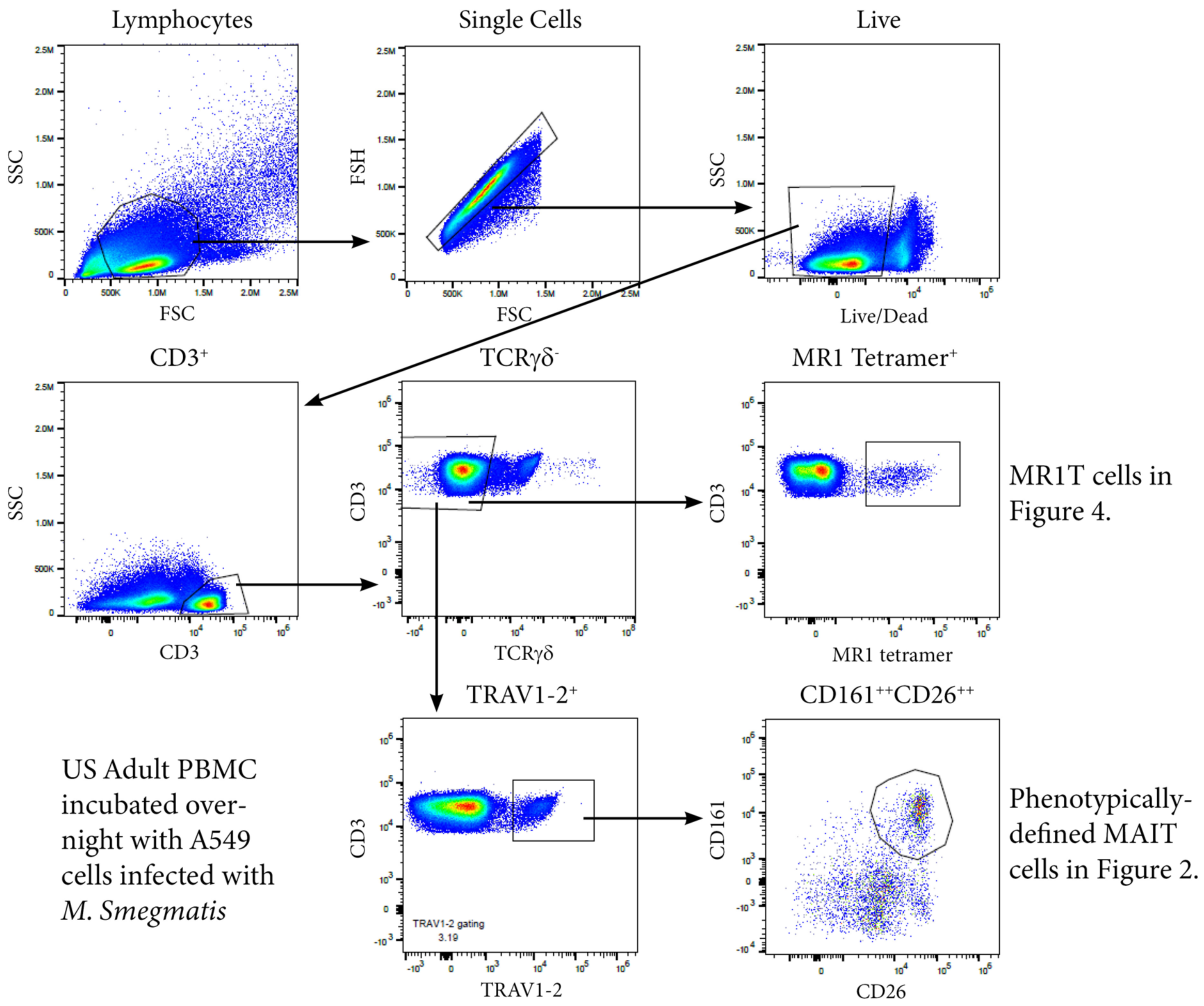


**Figure S6. Gating strategy for functional MR1T cells and phenotypically-defined MAIT cells.** PBMC or CBMC from the US cohort were incubated overnight with *M. smegmatis*-infected A549 cells, uninfected A549 cells, or uninfected A549 cells and PMA/ionomycin. All cells were then stained with the MR1-5-OP-RU or MR1-6FP tetramers, followed by a live/dead discriminator and antibodies to TCRγδ, CD3, CD4, CD8, TRAV1-2, CD26 and CD161. ICS was then performed and the cells stained for TNF. Live, TCRγδ$^-$CD3$^+$TRAV1-2$^+$CD161$^{++}$CD26$^{++}$ cells or live, TCRγδ$^-$ CD3$^+$MR1-5-OP-RU$^+$ cells were gated. Shown in the example is PBMC from an adult incubated overnight with *M. Smegmatis* infected A549s.

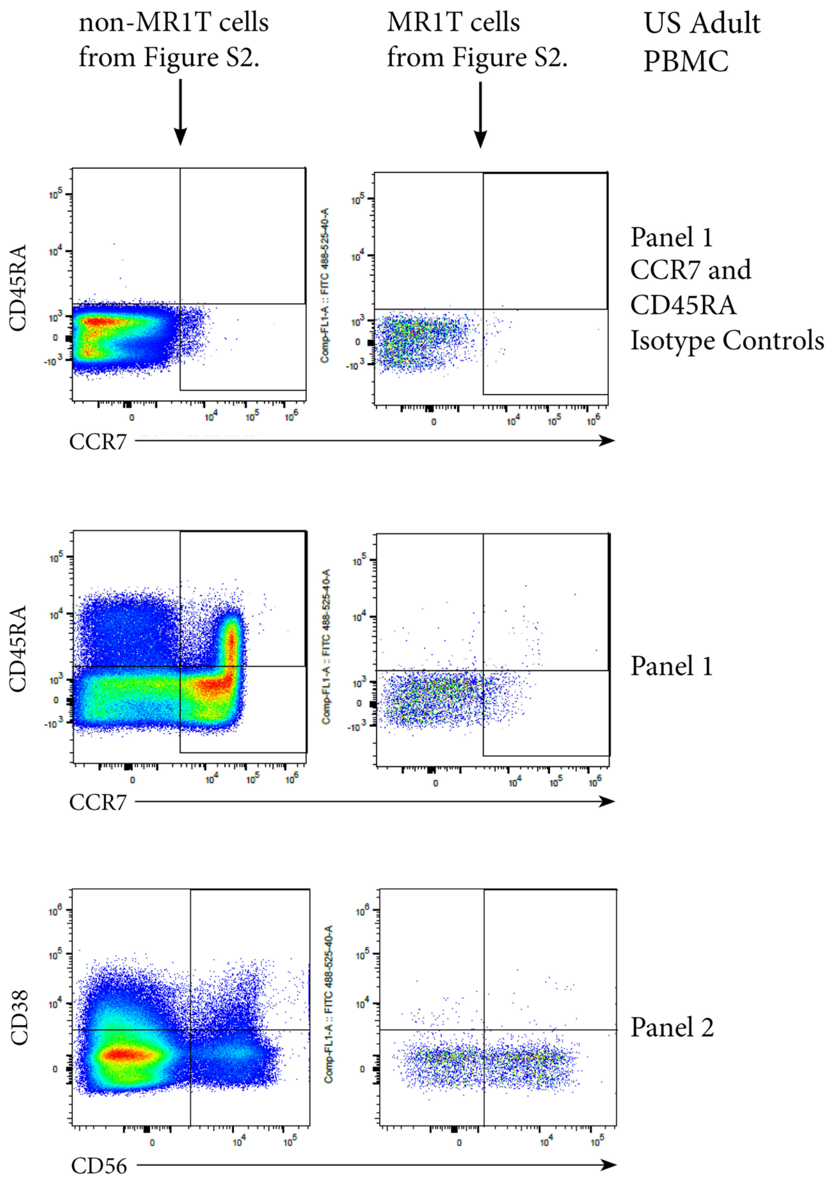


**Figure S7. Example of CD45RA, CCR7, CD38 and CD56 staining on MR1T and non-MR1T cells.** PBMC or CBMC from the US were stained as in Figure 1 and Supplementary Figure 3 with the addition of antibodies to either CD45RA, CCR7, CD38 or CD56. Also shown are the isotype controls for CD45RA and CCR7. Shown in the example is PBMC from an adult in the US cohort.

**Supplemental Table S1. Antibodies used for ICS assay in Figures 2 and 4.**

| Antibody | Flourochrome | Clone | Supplier |
|---|---|---|---|
| CD3 | FITC | OKT3 | Biolegend |
| CD26 | PerCP-Cy5.5 | BA5b | Biolegend |
| CD161 | PE-Cy7 | HP-3G10 | Biolegend |
| TCR gamma delta | APC | B1 | Biolegend |
| CD8 | APC-Cy7 | SK1 | Biolegend |
| CD4 | Brilliant Violet 785 | OKT4 | Biolegend |
| TRAV1-2 | Brilliant Violet 421 | 3C10 | Biolegend |
| TNF-α | Brilliant Violet 650 | Mab11 | Biolegend |
| Live/dead | Aqua | | ThermoFisher |
| MR1-5-OP-RU | PE | | NIH tetramer facility |
| MR1-6-FP | PE | | NIH tetramer facility |

**Table S2. Antibodies used for flow cytometry (US cohorts) for Figures 1, 3, 5 and 6.**

Three panels were run, identical except for different markers in FITC and APC as shown below.

| Antibody | Flourochrome | Clone | Supplier |
|---|---|---|---|
| CD3 | Brilliant Violet 650 | OKT3 | Biolegend |
| CD26 | PerCP-Cy5.5 | BA5b | Biolegend |
| CD161 | PE-Cy7 | HP-3G10 | Biolegend |
| CD8 | APC-Cy7 | SK1 | Biolegend |
| CD4 | Brilliant Violet 785 | OKT4 | Biolegend |
| TRAV1-2 | Brilliant Violet 421 | 3C10 | Biolegend |
| CD45RA | FITC (panel 1) | HP-3G10 | Biolegend |
| CD38 | FITC (panel 2) | HIT2 | Biolegend |
| CCR7 | APC (panel 1) | G043H7 | Biolegend |
| CD56 | APC (panel 2) | My51 | Tonbo |
| CD27 | APC (panel 3) | O323 | Biolegend |
| Live/dead | Aqua | | ThermoFisher |
| MR1-5-OP-RU | PE | | NIH tetramer facility |
| MR1-6-FP | PE | | NIH tetramer facility |

**S3 Table. Antibodies used for flow cytometry (South African Cohorts) for Figures 1 and 3.**

| Antibody | Flourochrome | Clone | Supplier |
|---|---|---|---|
| Live/Dead | Near Infrared | | Thermofischer |
| CD14 | APC-H7 | MAB | BD |
| CD19 | APC-H7 | SJ25C1 | BD |
| CD3 | Alexa Fluor 700 | UCHT1 | Biolegend |
| CD4 | Brilliant Violet 510 | RPA-T4 | Biolegend |
| CD8 | Brilliant Violet 785 | SK1 | Biolegend |
| CD26 | Brilliant Violet 605 | M-A261 | BD |
| CD161 | PE-Cy5 | DX12 | BD |
| TRAV1-2 | PE-Cy7 | 3C10 | Biolegend |
| MR1-5-OP-RU | PE | | NIH tetramer facility |
| MR1-6-FP | Alexa Fluor 488 | | NIH tetramer facility |

**S4 Table. Antibodies used for flow cytometry (Ugandan Cohorts) for Figures 1 and Supplementary Figure 1.**

| Antibody | Flourochrome | Clone | Supplier |
|---|---|---|---|
| CD3 | Brilliant Violet 650 | OKT3 | Biolegend |
| CD26 | FITC | BA5b | Biolegend |
| CD161 | PE-Cy7 | HP-3G10 | Biolegend |
| CD8 | APC-Cy7 | SK1 | Biolegend |
| CD4 | Brilliant Violet 421 | OKT4 | Biolegend |
| TRAV1-2 | APC | OF5A12 | Biolegend |
| Live/dead | Aqua | | ThermoFisher |
| MR1-5-OP-RU | PE | | NIH tetramer facility |
| MR1-6-FP | PE | | NIH tetramer facility |
